# Visual field position shapes input sampling and output routing in the superior colliculus

**DOI:** 10.64898/2026.05.27.728073

**Authors:** Alex Calzoni, Arnau Sans-Dublanc, Norma K. Kühn, Daan Remans, Fernando Fernandez de Cuevas Lopez, Katja Reinhard, Karl Farrow

**Affiliations:** VIB – KU Leuven Center of Neuroscience, Leuven, Belgium; Dept of Biology and Leuven Brain Institute, KU Leuven, Leuven, Belgium; ISTA, Austria; SISSA, Trieste, Italy

## Abstract

Animals use the location of visual stimuli to select appropriate actions^1–5^, and the upper and lower visual field often carry different ecological and behavioral meaning^6–9^. In mice, the superior colliculus is a key central hub that transforms visual input into orienting, defensive, and approach behaviors^3,10–13^. Its superficial layers receive retinotopically organized input from the retina and contain genetically defined cell types with distinct downstream projections, including wide-field neurons that project to the lateral posterior thalamus and narrow-field neurons that target the parabigeminal nucleus and deeper collicular layers^14–18^. These features raise the question of whether circuits of the superior colliculus are repeated across visual space or exhibit visual-field-dependent specializations. Here, we show that the mouse superficial superior colliculus contains visual-field-dependent circuit modules. Dual-color rabies tracing revealed that wide-field and narrow-field neurons receive input from a largely shared set of brain regions, whereas upper- and lower-field domains differ in how they sample those inputs. Some source regions preferentially innervate one visual-field domain, producing biased regional input strength, while others contain topographically segregated projecting neurons that target upper- or lower-field domains. MAPseq showed that most superficial collicular neurons project to single downstream targets, with upper- and lower-field populations differing in target probability. Two-photon calcium imaging further showed that wide-field neurons in upper- and lower-field domains differ in stimulus selectivity. Together, these findings reveal a visual-field-dependent wiring logic that biases how the superior colliculus samples inputs and routes signals to downstream pathways.

**Highlights:**

- Wide- and narrow-field neurons receive broadly overlapping inputs
- Upper- and lower-field domains differ in input strength and topographic organization
- Most superficial collicular neurons project to a single target
- Visual field position biases downstream target probability

## Results

### Wide- and narrow-field neurons sample a shared set of brain areas

To compare brain-wide inputs to upper- and lower-field neurons in the superficial layers of the superior colliculus (SCs, referred to in text as ‘colliculus’), we used dual-color monosynaptic rabies tracing in Ntsr1-GN209-Cre and Grp-KH288-Cre mice^19^ to target wide-field and narrow-field neurons, respectively. Cre-dependent helper viruses (HSV-FLEX-TVA-N2c(G)-mCherry for Ntsr1 mice and AAV9 -FLEX-TVA-N2c(G)-mTagBFP2 for Grp mice) were delivered to the SCs, followed by EnvA-pseudo-typed rabies viruses expressing different fluorescent proteins (CVS-RV-EnvA-GFP and CVS-RV-EnvA-tdTomato) injected into medial and lateral SCs, corresponding to upper- and lower-field (Fig 1a). We verified separation of the two domains in both the colliculus, and retina and registered labeled presynaptic neurons to the Allen Common Coordinate Framework for whole-brain quantification (Fig. 1c)^20,21^. Wide- and narrow-field neurons received inputs from a largely shared set of brain areas. Pooling dual-color rabies datasets within each animal identified 23 major input regions for each population (wide-field: 77,263 inputs, n = 8; narrow-field: 47,897 inputs, n = 7; proportion threshold ≥ 1%; see Methods). Log-transformed input fractions were strongly correlated across areas (Pearson’s r = 0.89; Fig. 2b), and the spatial distributions of input neurons along the medial-lateral, anterior-posterior, and dorsal-ventral axes were also similar (Fig. 2d). This similarity was supported by hierarchical clustering of input areas, which yielded comparable organization across the two cell types (Fig. S1d).

**Fig. 1.**
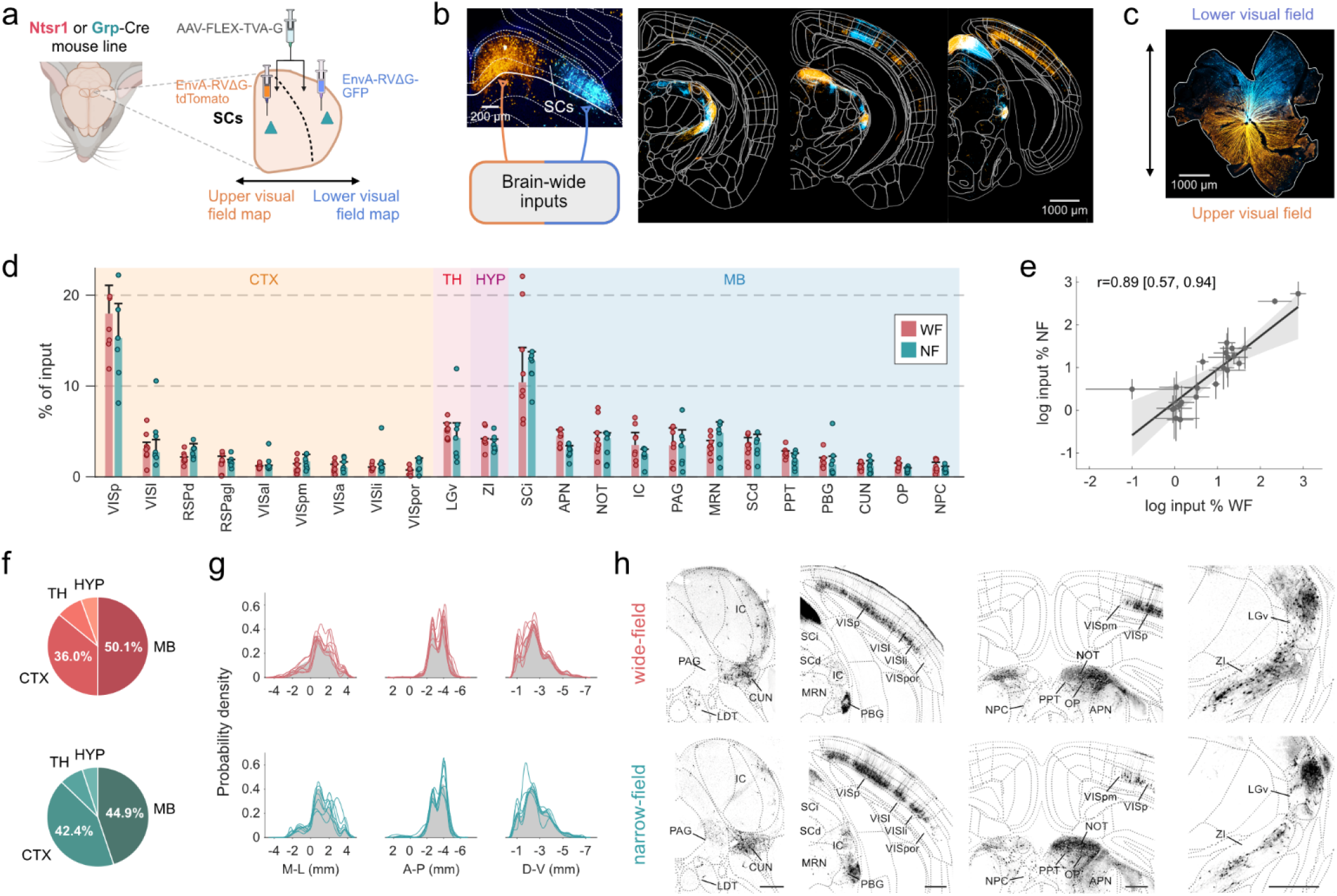
Wide- and narrow-field neurons sample a shared set of brain areas. **a** Schematic of the experimental approach. Cre-dependent helper virus (AAV-FLEX-TVA-G) was injected into the superficial layers of the colliculus (SCs) of Ntsr1-Cre or Grp-Cre mice (n = 7 upper, n = 7 lower in Ntsr1-Cre; n = 6 upper, n = 5 lower in Grp-Cre), followed by EnvA-pseudotyped, RG-deleted rabies viruses expressing tdTomato (medial SCs, upper visual field) or GFP (lateral SCs, lower visual field). **b** Coronal sections showing the SCs injection site and brain-wide distribution of labeled presynaptic neurons projecting to the two visual field domains. Presynaptic neurons of each color were counted separately and aligned to the Allen Brain Atlas. **c** Example retina showing spatial segregation of inputs from upper and lower visual field. **d** Percentage of total inputs from each brain area for wide-field (WF) and narrow-field (NF) neurons. Areas are grouped into cortex (CTX), thalamus (TH), hypothalamus (HYP), and midbrain (MB). Bars show median ± median absolute deviation (MAD); dots represent individual animals (wide-field: 77,263 inputs, n=8; narrow-field: 47,897 inputs, n=7). **e** Correlation of log-transformed input percentages between wide- and narrow-field inputs. Each point represents one brain area; error bars show MAD. Shading indicates 95% bootstrap confidence interval. **f** Proportion of inputs by major brain division for wide-field (red) and narrow-field (teal) neurons. **g** Probability density distributions of input neurons along medial-lateral (M-L), anterior-posterior (A-P), and dorsal-ventral (D-V) axes. Gray shading shows the mean density distribution, colored lines represent individual animals. **h** Example coronal sections labeling input neurons across midbrain, visual cortex, and thalamic regions. Scale bars, 500 µm.

**Fig. 2.**
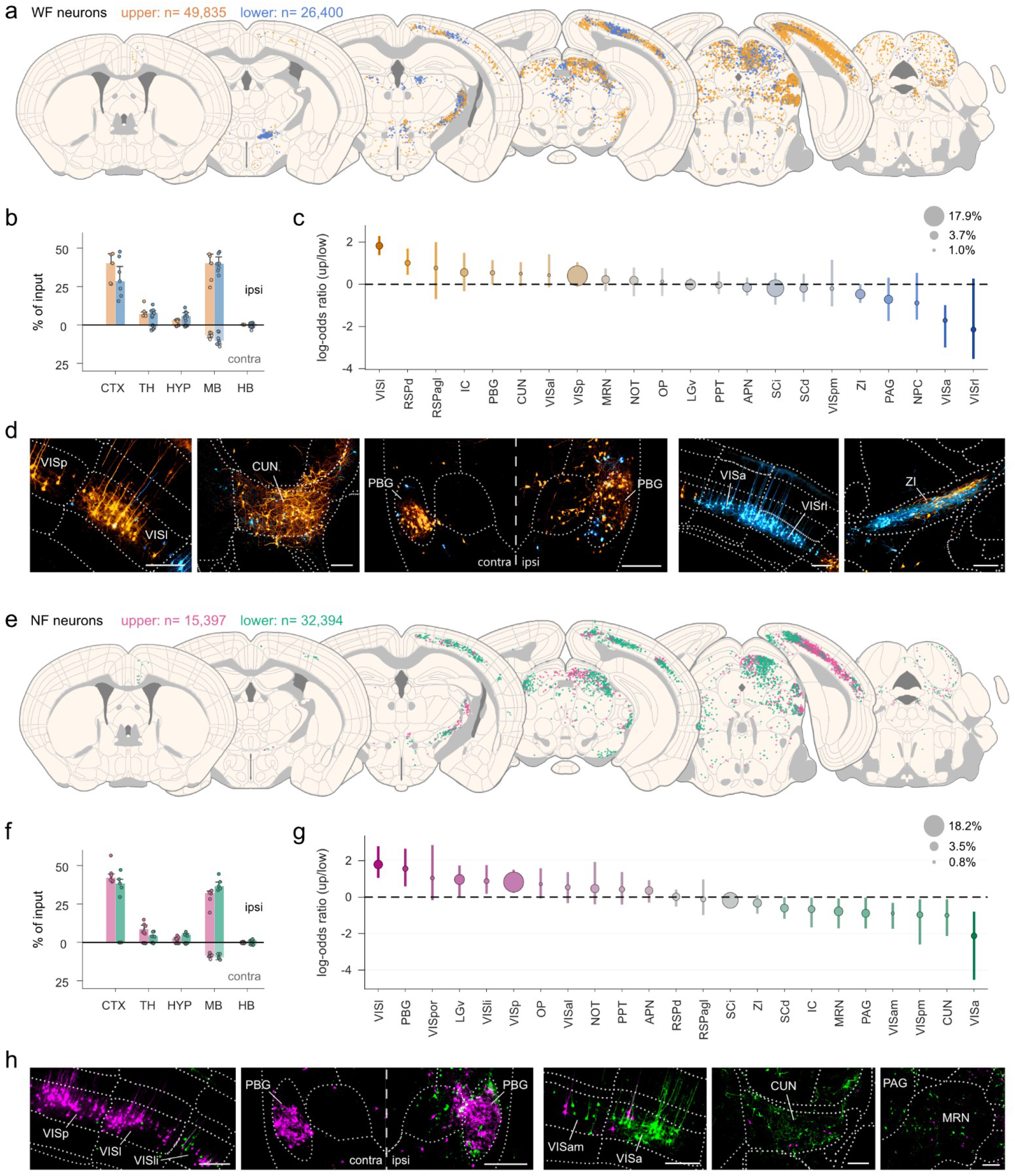
Preferential inputs to upper- and lower-visual-field domains. **a** Coronal sections with input neurons projecting to upper (49,835 inputs n=7) and lower (26,400 inputs, n=7) visual field wide-field neurons. Each section displays neurons within 100 μm, pooled across animals. **b** Percentage of total inputs by major brain division for ipsilateral and contralateral hemispheres. Bars show median ± median absolute deviation; dots represent individual animals. **c** Log-odds ratios comparing input proportions between visual field domains, sorted by effect size. Positive values indicate upper visual field bias; negative values indicate lower visual field bias. Error bars show 95% bootstrap confidence intervals corrected for multiple comparisons (false coverage rate). Circle size reflects mean input percentage. **d** Example coronal sections showing input neurons in regions with domain-specific biases. **e–h** Same as a–d for narrow-field neurons (15,397 inputs, n=6 for upper; 32,394 inputs, n=5 for lower visual field). Throughout, orange/blue denote upper/lower visual field for wide-field neurons; magenta/green denote upper/lower visual field for narrow-field neurons.

To test for cell-type-specific biases, we computed log-odds ratios comparing input fractions between wide-field and narrow-field neurons (see Methods). The anterior pretectal nucleus (APN) was the only region showing an enrichment for wide-field neurons, whereas all other major input areas showed no detectable cell-type bias (Fig. S1e). In contrast, Gad2-expressing inhibitory neurons, another major cell type in the SCs^17^, received proportionally more hypothalamic and hindbrain input and less midbrain input than both wide- and narrow-field neurons (Fig. S1f-h). Thus, despite their distinct downstream targets and behavioral associations, wide- and narrow-field neurons sample broadly overlapping upstream networks.

### Certain input areas preferentially target upper or lower visual field neurons

While wide- and narrow-field neurons shared a common input architecture, a subset of brain regions preferentially targeted upper- or lower-field domains of the colliculus in both cell types. To quantify these biases, we computed log-odds ratios comparing the fraction of inputs to upper- and lower-field domains, where positive values indicate upper-field bias and negative values indicate lower-field bias (Fig. 2c,e). In wide-field neurons, the bias estimates formed a gradient from regions favoring upper-field domains, including VISl and RSPd, with weaker trends in PBG and CUN, to regions favoring lower-field domains, including VISa and ZI (Fig. 2d). Narrow-field neurons showed biases in many of the same areas, including VISl, VISa, and PBG. CUN was also biased in both cell types but in opposite directions. Narrow-field neurons additionally showed biases in several visual cortical areas (VISp, VISli, VISpm, VISam) and in MRN (Fig. 2h). These cortical biases are consistent with the functional retinotopic maps reported by Zhuang et al. (2017)^22^, where higher-order areas such as VISl and VISa show biased representations of visual space. These domain-specific biases occurred against a background of broadly shared input architecture. Across both cell types, inputs arose predominantly from ipsilateral cortex and midbrain, with smaller contributions from thalamus, hypothalamus, and contralateral midbrain (Fig. 2a,b,e,f). Major brain divisions, as well as the midbrain subdivisions MBmot, MBsen, and MBsta, contributed similar fractions of total input to upper- and lower-field domains (Fig. S2a). The overall pattern of domain bias was broadly similar across cell types, but differed in specific regions. Correlations between the log-transformed upper- and lower-field input fractions were strong for wide-field neurons (r = 0.74) and more moderate for narrow-field neurons (r = 0.55), indicating that most regions contributed similarly to both domains, but that deviations were larger in narrow-field circuits (Fig. S2g). Across all areas, log-odds estimates for wide-field and narrow-field neurons were moderately correlated (r = 0.66; Fig. S2h). To test whether these domain biases differed between cell types, we computed interaction effects as the difference in log-odds ratio between wide-field and narrow-field neurons. Regions including CUN, IC, MRN, and RSPd showed interaction estimates whose confidence intervals excluded zero, indicating potential cell-type-dependent differences in domain bias (Fig. S2e).

### Several areas show topographic organization of inputs

Differences in input proportions are not the only way retinotopic specificity could arise. Neurons projecting to upper- and lower-field domains may also be spatially segregated within the source regions themselves. We therefore identified, for each area and cell type, the spatial axis that maximally separated neurons projecting to upper-versus lower-field domains (see Methods; Fig. S3a). We computed kernel density estimates along candidate directions in 3D space and selected the axis that minimized distribution overlap. Separation was expressed as a fraction ranging from 0 (complete overlap) to 1 (complete segregation), and was averaged across animals for each region (Fig. 3a; Fig. S3d,e).

**Fig. 3.**
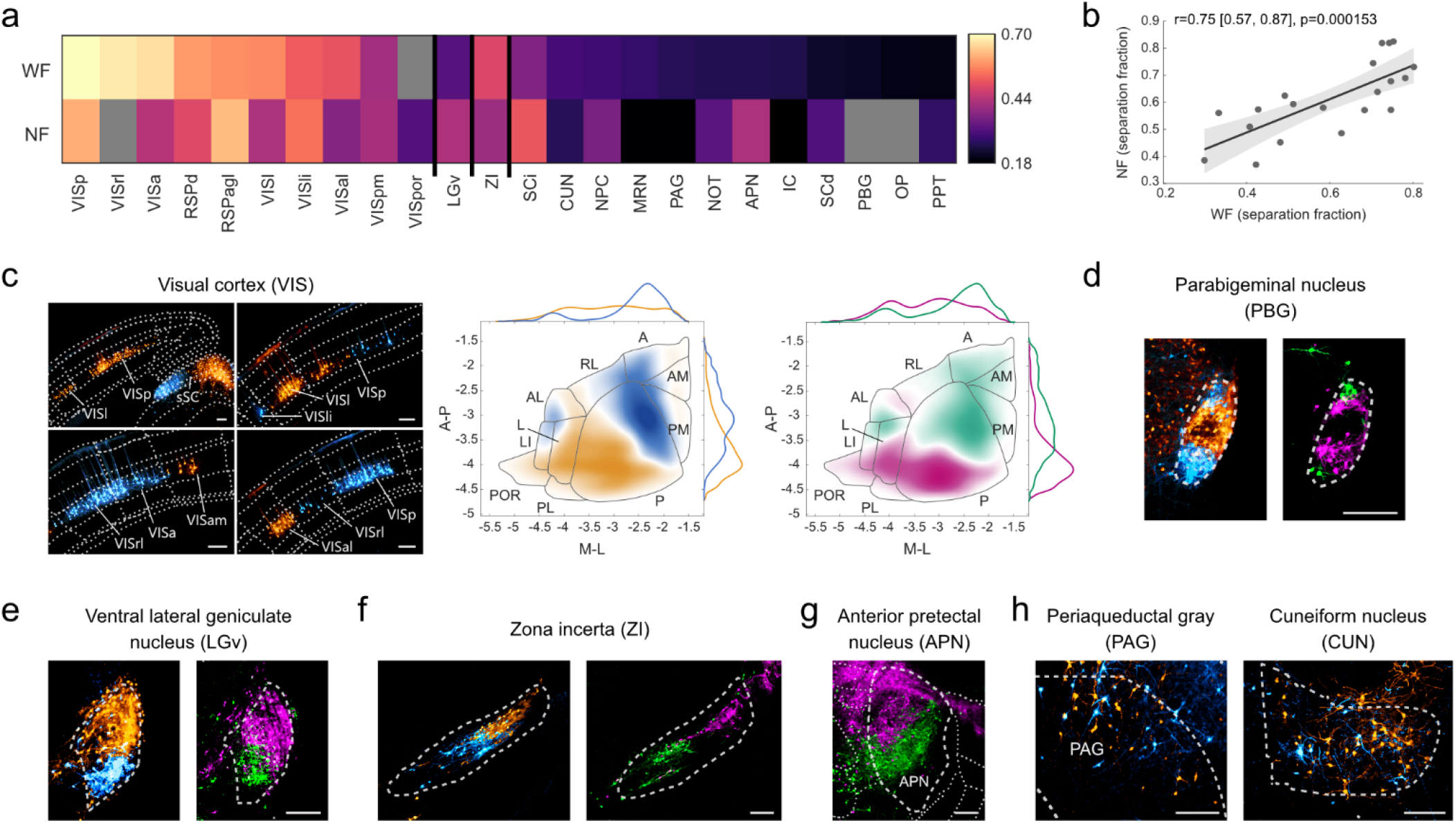
Topographic organization of inputs. **a** Spatial segregation of neurons projecting to upper versus lower visual field domains within each input region. Separation fraction ranges from 0 (complete overlap) to 1 (complete segregation), computed along the axis of maximum separation and averaged across animals. Gray indicates insufficient neurons for analysis. **b** Correlation of separation fractions between wide-field and narrow-field neurons. Each point represents one brain area. Shading indicates 95% bootstrap confidence interval. **c** Topographic organization in visual cortex. Left: coronal sections with input neurons to wide-field neurons. Right: kernel density estimates projected onto visual cortical areas for wide-field and narrow-field neurons. **d-g** Coronal sections illustrating topographic segregation of upper and lower visual field-projecting neurons in parabigeminal nucleus (d), ventral lateral geniculate nucleus (e), zona incerta (f), and anterior pretectal nucleus (g). **h** Example regions with no apparent topographic segregation. Throughout, orange/blue denote upper/lower visual field for wide-field neurons; magenta/green denote upper/lower visual field for narrow-field neurons. Scale bars, 200 μm.

Several cortical and subcortical regions contained spatially segregated populations of neurons projecting to upper- and lower-field SCs domains. In cortex, visual and retrosplenial areas showed clear topographic alignment, where neurons in regions known to represent the upper or lower visual field^22^ projected to the corresponding SCs domain (Fig. 3c; Fig. S3b). Similar segregation was evident in several subcortical structures, including PBG, LGv, ZI, APN, and NPC (Fig. 3d-g; Fig. S3c). Separation estimates were correlated across the two cell types (r = 0.75; Fig. 3b), although segregation was more pronounced for narrow-field inputs in several midbrain regions. Some areas showed clear topographic segregation despite little or no bias in input proportion, whereas others showed domain bias without obvious spatial segregation (e.g. CUN, Fig. 3h). These results indicate that retinotopic specificity in SC input sampling can arise through two distinct mechanisms: differences in the amount of input contributed by a source region and spatial segregation of the neurons within that region that project to upper- and lower-field domains.

### MAPseq reveals single-target projections that differ across visual field domains

Having identified retinotopic biases in input sampling, we next asked whether upper- and lower-field populations also differ in how they route signals to downstream targets. We employed Multiplexed Analysis of Projections by Sequencing (MAPseq), a high-throughput single-neuron tracing technique that uniquely labels thousands of neurons with randomized RNA barcodes23. Barcodes were sequenced after after dissecting five target areas (LP, PBG, LGd, LGv, and the mesencephalic locomotor region, MLR24) from both hemispheres, along with source regions in the medial and lateral SCs corresponding to the upper and lower visual fields (Fig. 4a). The relative barcode abundance in after dissecting five target areas (LP, PBG, LGd, LGv, and the mesencephalic locomotor region, MLR24) from both hemispheres, along with source regions in the medial and lateral SCs corresponding to the upper and lower visual fields (Fig. 4a). The relative barcode abundance in each target area served as a measure of projection strength (density of axonal endings). Most labeled neurons projected to only one of the five assayed targets, with 93.6% of upper (288/307) and 89.7% of lower (240/267) SCs neurons exhibiting single-target specificity, ∼4.8% and 9.2% bifurcating to two targets, and ∼1.6% and 1.1% projecting to three or more targets (Fig. 4b,c). LP was the most common projection target for both populations, receiving projections from 58% of upper and 44% of lower field neurons (Fig. 4b and S4b). The fraction of dedicated outputs varied by target region, with PBG-projecting neurons showing the highest fraction of single projections (97.6% and 100% for upper and lower SCs), while upper SCs neurons projecting to the LGv had the lowest at 40% (Fig. 4d). The vast majority of projections were ipsilateral (95.9% of upper and 97.1% of lower SCs neurons, Fig. S4c).

**Fig. 4.**
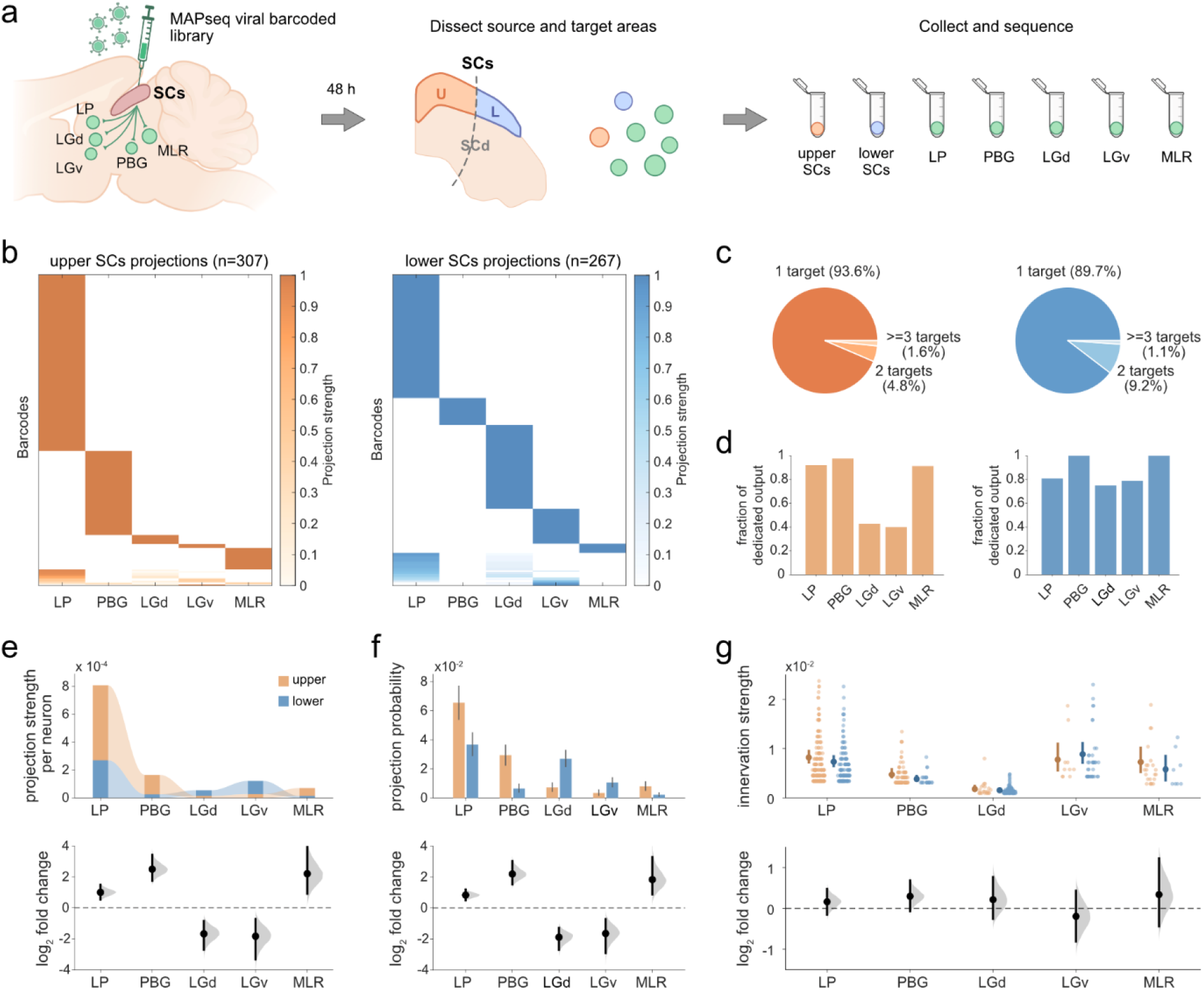
MAPseq reveals single-target projections that differ across visual field domains. **a** Schematic of MAPseq approach. Barcoded Sindbis virus was injected into the SCs, labeling neurons with unique RNA barcodes. After 48 hours, source regions (upper and lower SCs) and target areas (LP, PBG, LGd, LGv, MLR) were dissected and sequenced. **b** Sorted heatmaps of relative projection strength of individual neurons from upper (n = 307) and lower (n = 267) SCs to the five targets. **c** Proportion of neurons projecting to one, two, or three or more targets for upper and lower SCs. **d** Fraction of dedicated output (single-target projections) for each target area. **e** Average projection strength per neuron (top) and log_2_ fold-change between upper and lower SCs (bottom). **f** Projection probability (top) and log_2_ fold-change (bottom). **g** Innervation strength for neurons projecting to each target (top) and log_2_ fold-change (bottom). Error bars show 95% bootstrap confidence intervals corrected for multiple comparisons (false coverage rate). Throughout, orange denotes upper and blue denotes lower visual field SCs.

Upper- and lower-field populations differed in target probability, with upper field neurons projecting more strongly to LP, PBG, and MLR, and lower field neurons preferencing the LGd and LGv (Fig. 4e). These differences were driven primarily by projection probability rather than by innervation strength among projecting neurons. Upper field neurons were ∼1.8×, 4.6×, and 3.5× more likely to innervate LP, PBG, and MLR, whereas lower field neurons were ∼3.7× and ∼3× more likely to target LGd and LGv (Fig. 4f). Innervation strengths were similar across all targets for both populations (Fig. 4g). Projection biases were also observed along the anterior-posterior axis (corresponding to nasal and temporal visual field), with posterior SCs neurons preferentially targeting LP and MLR (Fig. S4e). Together, these results indicate that SCs output organization is dominated by single-target, ipsilateral projections, and suggest that upper and lower visual field, as well as anterior and posterior neuron populations can exhibit distinct target preferences.

Because a small fraction of neurons projected to more than one target, we tested whether these projection motifs deviated from expectations based on independent target probabilities. Most observed motifs were well explained by a binomial model, and deviations from independence were small in both upper- and lower-field populations (Fig. S4g-l). The main exception was an underrepresentation of LP-PBG bifurcation among upper-field neurons, consistent with the high fraction of dedicated PBG-projecting neurons (Fig. 4d). Beyond long-range targets, local SCs projections preserved retinotopic alignment, with each superficial subdivision preferentially targeting its corresponding deeper subdivision (Fig. S4n).

Additionally, using two-photon calcium imaging, we found that wide-field neurons in upper- and lower-field domains differed in stimulus selectivity: upper-field neurons responded more selectively to black looming discs, whereas lower-field neurons responded to a broader range of stimulus classes (Fig. S5).

Disynaptic tracing through LP further suggested partially distinct downstream routing, with labeling in caudoputamen and amygdala observed only for the upper-field condition, though this difference should be interpreted cautiously given limited sample size (Fig. S6).

## Discussion

We asked whether the connectivity of SCs cell types differs across visual field domains. Three conclusions emerge. First, wide-field and narrow-field neurons receive input from a largely shared set of ∼23 cortical, thalamic, hypothalamic, and midbrain regions. Second, within that shared landscape, upper- and lower-field representations differ in two ways: some source regions are biased in the proportion of input they contribute to each domain, and other regions contain spatially segregated populations of neurons that project to upper-versus lower-field representations. Third, neurons in the SCs project predominantly to single downstream targets, and upper- and lower-field populations differ in target probability. Together, these results support a modular organization in which visual-field position biases both input sampling and output routing in the colliculus.

The broad similarity of inputs to wide-field and narrow-field neurons was not expected. Previous work emphasized differences in their output targets and role in behavior^25,14,17^, which suggested that they might also sample different presynaptic partners. Instead, wide- and narrow-field neurons sampled from a common set of brain areas. Previous studies in mice and other rodents have identified the major inputs to the SCs from retina, visual and retrosplenial cortices, LP, LGd, LGv, PBG and pretectum^26–30^. Additionally, our results show that both wide- and narrow-field neurons receive inputs from regions including the ZI, IC, MRN, PAG, PPT and CUN, consistent with previous findings from our laboratory in Gad2 neurons^31^. These regions may have gone undetected in earlier work because most studies used conventional retrograde tracers, with few employing rabies vectors^26^, and our use of the CVS rabies strain, which exhibits high transsynaptic efficiency^32^, likely increased sensitivity to sparse projections.

Despite this similarity, our input mapping reveals two forms of visual-field-specific input selectivity. Some regions differed in the fraction of inputs sampled by upper- and lower-field domains, whereas others showed topographic segregation of projecting neurons without a large difference in total input fraction. Topographic segregation in visual and retrosplenial cortices formed a continuous map, consistent with previous reports that retinotopy extends from visual cortex into retrosplenial areas^22^. These two patterns imply distinct wiring rules. A biased input fraction changes how much influence a source region can exert on one domain relative to the other. Topographic segregation instead preserves visual-field specificity. These two forms of selectivity were independent: some areas showed domain preference without topographic segregation, while others showed segregation without input bias. The presence of both motifs argues against a simple model in which the SCs consists of one repeated circuit tiled across retinotopic space.

The output data extend this view from input structure to routing. As the first application of MAPseq to the colliculus, MAPseq showed that most SCs neurons in our dataset projected to a single target and that upper- and lower-field populations differed primarily in projection probability rather than in innervation strength among projecting neurons. That is, the probability of entering a given pathway varies with retinotopic location, not the density of the axonal endings. Upper-field neurons were biased toward parabigeminal and locomotor-related targets, whereas lower-field neurons were biased toward targets in the thalamus. The few multi-target motifs observed had low counts, and most were explained by a binomial model, suggesting bifurcations arise stochastically rather than from structured rules. One exception was the underrepresented LP-PBG bifurcation, consistent with previous findings that LP-projecting and PBG-projecting neurons form distinct dedicated pathways^31,17^.

Consistent with this circuit asymmetry, wide-field neurons in upper- and lower-field domains differed in stimulus selectivity, and disynaptic tracing through LP suggested partially distinct downstream targets (Figs. S5, S6). These findings raise the possibility that retinotopic modules differ not only in how they route signals but also in the visual features to which they respond. In mice, overhead stimuli in the upper visual field typically trigger defensive responses such as freezing or escape, while stimuli in the lower field more often elicit approach^2,6,7,33,34^. The visual-field-dependent differences in connectivity and stimulus selectivity reported here may contribute to these behavioral asymmetries, consistent with prior evidence that upper-field circuits are important for defensive responses to overhead stimuli^3,16,34^.

Several limitations should be considered when interpreting these findings. Input mapping may be affected by atlas-registration errors, threshold choice, and variation in injection depth, especially in the lateral colliculus where superficial layers are thin. The Cre lines used here also label neurons outside the intended superficial populations^14^. MAPseq was performed in three mice, so analysis was fixed-effect at the neuron level^35^. Passing fibers may have inflated apparent multi-target motifs, especially among adjacent regions such as LP, LGd, and LGv. These caveats do not alter the main conclusion that upper- and lower-field populations differ in connectivity, but they do limit the precision with which individual regional effects can be estimated.

In summary, the superficial colliculus is organized into visual-field defined modules with biased input sampling and biased output routing. This organization provides a circuit framework through which identical visual stimuli can engage different pathways depending on their location in the visual field. A next step will be to link these modules more directly to molecularly defined cell classes and to the behavioral computations they support.

## Acknowledgements

We thank Chen Li for setting up the SHARP-track toolbox for brain atlas registration and Frédérique Ooms for help with histology and immunohistochemistry. Grants: FWO fellowship 1159422N/1159424N (A.C.), FWO fellowship 1197818N/1197820N (A.S.-D.), FWO fellowship 1205421N (N.K.K.), FWO fellowship 12S7917N/ 12S7920N (K.R.), FWO grant G091719N (K.F.), NIH grant 1RO1EY032101 (K.F.), and KU Leuven Research Council grant C14_22_074 (K.F.).

## Author contributions

A.C. and K.F. conceived and designed the study. A.C. and A.S.-D. performed the rabies tracing experiments. A.C. carried out the cell counting, atlas registration, and all associated analyses. K.R. and A.S.-D. performed the MAPseq injections and tissue dissections; A.C. analyzed the MAPseq data. N.K.K. and D.R performed the calcium imaging experiments and analysis of wide-field neuron response properties. N.K.K. and F.F.d.C.L. performed the disynaptic tracing experiments and histological analysis. A.C. and N.K.K. generated the figures. A.C., N.K.K., and K.F wrote the manuscript.

## Star Methods

### Key resources table

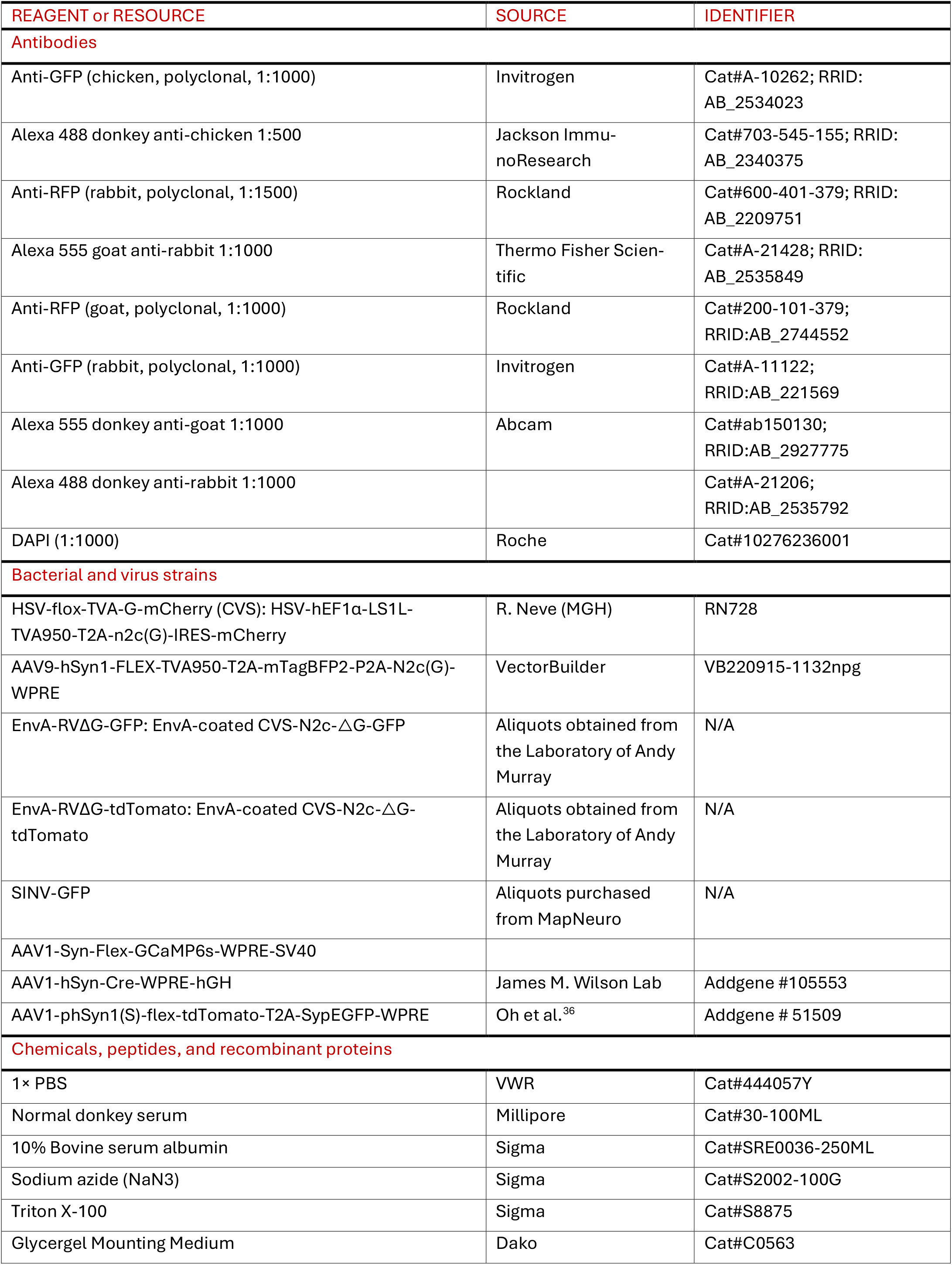
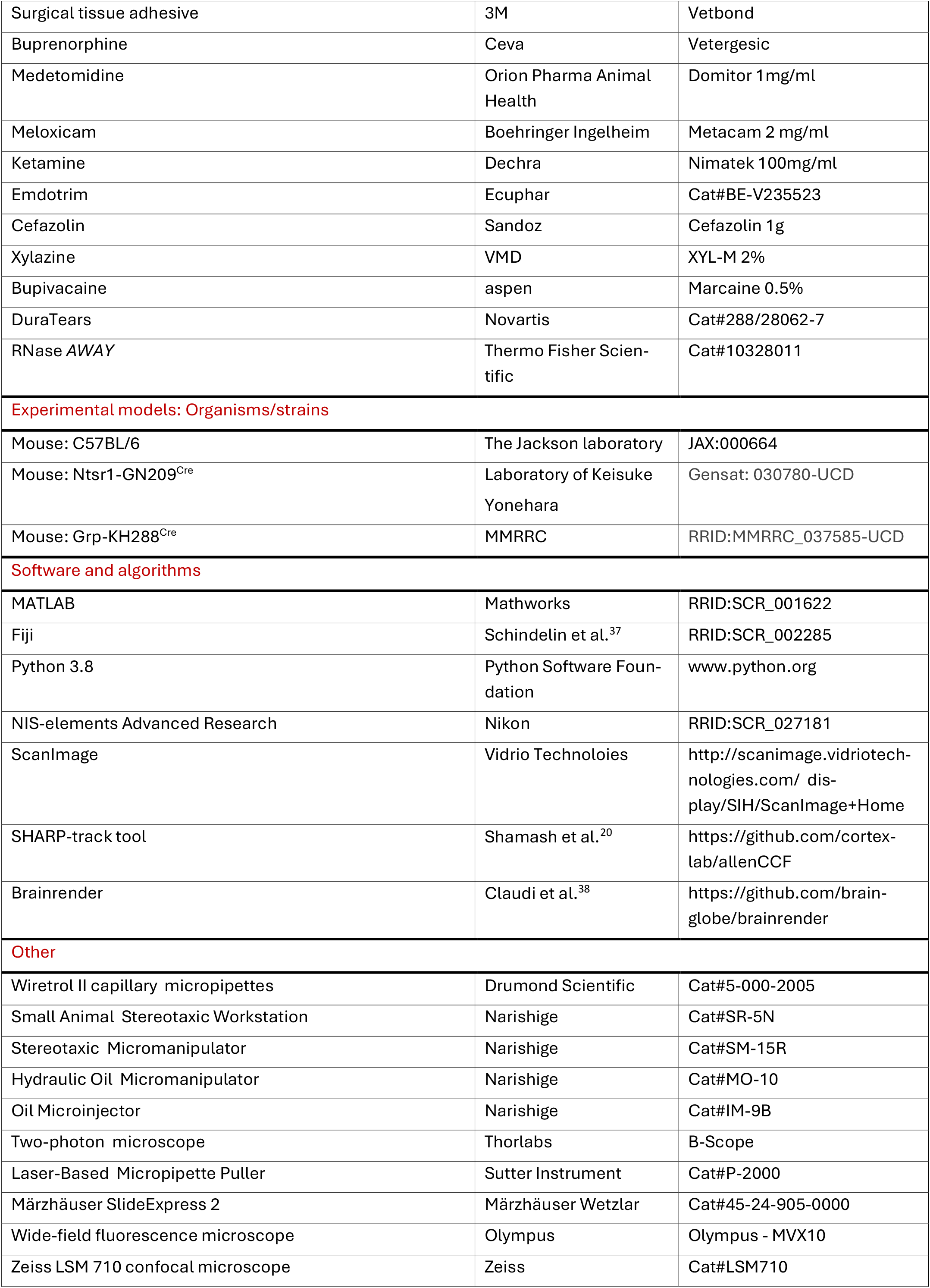

### Contact for reagent and resource sharing

Further information and requests for resources and reagents should be directed to and will be fulfilled by Karl Farrow.

### Animals

All experimental procedures were approved by the Ethical Committee for Animal Experimentation (ECD) of the KU Leuven and followed the European Communities Guidelines on the Care and Use of Laboratory Animals (014-2018/EEC). Male and female adult (2-7 months) transgenic (Ntsr1-GN209-Cre and Grp-KH288-Cre^19^) and WT (C57Bl/6J) mice were used. Mice were housed under a 12h light-dark cycle (lights on at 7:00 a.m.) and provided with food pellets and water ad libitum. We used 8 Ntsr1-Cre mice and 7 Grp-Cre mice for brain-wide input tracing, 3 WT mice for MAPseq, 7 Ntsr1-Cre mice for two-photon calcium imaging, and 5 Ntsr1-Cre mice for disynaptic tracing.

### Stereotactic surgeries

Mice were treated preoperatively with buprenorphine (0.4 mg/kg subcutaneously; Vetergesic). Anesthesia was induced with a combination of Ketamine (75mg/kg) and Medetomidine (1mg/kg) injected subcutaneously. Mice were placed in a stereotaxic workstation (Narishige, SR-5N) and kept at physiological temperature by a homeothermic blanket. After achieving deep anesthesia, dura tear (NOVARTIS, 288/28062-7) was applied to protect the eyes from drying out. The scalp was shaved, disinfected (Iso-Betadine), and the local anesthetic Bupivacaine (0.5%, 150 µl) was injected locally. After five minutes, the skull was exposed with a single incision and cleaned with saline. The location of the medial and lateral SCs was determined based on bregma and lambda. Holes were created at the measured coordinates (see below) by gently rotating a needle against the skull. We used micropipettes (Wiretrol II capillary micropipettes, Drumond Scientific, 5-000-2005) with an open tip of approximately 30 μm and an oil-based hydraulic micromanipulator MO-10 (Narishige) for virus injections. After the injection, the micropipette was slowly removed; the skull was cleaned with Betadine and the scalp was closed using Vetbond tissue adhesive (3M, 1469). After surgery, mice were allowed to recover on top of a heating pad, treated with analgesics (buprenorphine 0.4 mg/kg and meloxicam 5 mg/kg, diluted in saline), and provided with soft food and water containing antibiotics (emdotrim, ecuphar, BE-V235523) for 48 hours.

### Brain-wide input tracing

#### Rabies retrograde tracing injections

To trace brain-wide inputs to specific cell types of the colliculus, we used monosynaptic retrograde tracing based on deletion-mutant rabies virus (RV). The first injection transduced wide-or narrow-field neurons in the SCs with a Cre-dependent helper AAV expressing the rabies glycoprotein G and the receptor protein TVA, so that a subsequent injection with deletion-mutant rabies virus would trace inputs exclusively to G- and TVA-expressing neurons. Because Cre expression occurs across different layers of the SC and other brain nuclei in these Cre lines^14^, we aimed to minimize off-target effects by limiting expression to the superficial layers. For wide-field neurons, the only SCs neurons to innervate the LP and which exclusively project to this nucleus^17^, we injected a Cre-dependent retrograde HSV helper (HSV-flox-TVA-G-mCherry) into the left LP (coordinates: AP -2.0, ML -1.6, DV -2.5, based on a bregma-lambda distance of 4.7 mm; 150 nl at a rate of 50 nl/min). After injection, the micropipette was held in place for 10 minutes before slowly retracting it. This approach was not feasible for narrow-field neurons, which project to multiple output regions also targeted by other SCs cell types^17^. We thus injected a Cre-dependent AAV helper (AAV9-hSyn1-FLEX-TVA950-T2A-mTagBFP2-P2A-N2c(G)-WPRE) into the left medial and lateral SCs of Grp-Cre mice (coordinates: medial AP -3.7, ML -0.3, DV -1.25, -1.2; lateral AP -3.9, ML -1.1, DV -1.35, -1.3; based on a bregma-lambda distance of 4.21 mm; 25 nl per injection depth = 50 nl total at a rate of 25 nl/min, waiting 5 minutes between injections). AP positions were adjusted based on the bregma-lambda distance of each animal.

After 16-21 days, we injected EnvA-coated rabies viruses expressing different fluorescent proteins (CVS-RV-EnvA-GFP and CVS-RV-EnvA-tdTomato) in the left medial and lateral SCs of Ntsr1- and Grp-Cre mice (coordinates for bregma-lambda distance of 4.21 mm: medial AP -3.7, ML -0.3, DV -1.25, -1.2; lateral AP -3.9, ML -1.1, DV -1.35, -1.3; 40 nl per injection depth = 80 nl total at a rate of 25 nl/min, waiting 5 minutes between injections). We alternated injection of the two types of rabies viruses into the medial and lateral SCs between animals. Twelve days later, mice were given a lethal dose of anesthetic (ketamine 300 mg/kg & xylazine, 30 mg/kg) and transcardially perfused with 4% paraformaldehyde following loss of the pedal reflex. Two control mice (one Ntsr1-Cre, one Grp-Cre) were perfused 21 days after helper virus injection to assess Cre-dependent specificity. In both cases, labeled neurons were confined to the expected layers (SO for wide-field and SGS for narrow-field neurons) and displayed morphologies consistent with wide-field and narrow-field neurons.

#### Brain Immunohistochemistry

Extracted brains were post-fixed in 4% PFA at 4°C overnight. Coronal sections (100 μm) were collected in 1x PBS + 0.02% sodium azide using a vibratome. Sections were permeabilized in 1× PBST (0.3% Triton-X100 in PBS) for 20 minutes and blocked with PBST containing 10% normal donkey serum at 4°C overnight. Sections were then incubated with primary antibodies in PBST containing 3% normal donkey serum for 2-3 days at 4°C with shaking. Afterwards, sections were washed three times in PBST for 20 min each before incubation with secondary antibodies in PBST containing 3% normal donkey serum at 4°C overnight with shaking. Primary antibodies: rabbit anti-RFP (1:1500) and chicken anti-GFP (1:1000); secondary antibodies: goat anti-rabbit Alexa Fluor 555 (1:1000) and donkey anti-chicken Alexa Fluor 488 (1:500). Sections were washed three times in PBST for 15 min each and once in PBS for 10 minutes, then stained for nuclei with DAPI (1:1000) in PBS for 30 min at room temperature with shaking. Sections were mounted on glass slides using Dako Glycergel mounting medium.

#### Retina Immunohistochemistry

Dissected retinas were fixed in 4% PFA containing 100 mM sucrose for 30 min at 4°C, then washed in PBS + 0.02% sodium azide overnight at 4°C. Retinas were cryoprotected through sequential incubation in 10%, 20%, and 30% sucrose solutions (30 min, 1 h at room temperature, and overnight at 4°C, respectively). Freeze-cracking was performed by freezing retinas on slides covered with 30% sucrose on dry ice for 5 min, followed by thawing at room temperature; this cycle was repeated three times. Retinas were washed three times in PBS (10 min each), then blocked with 10% normal donkey serum, 1% BSA, 0.5% Triton X-100, and 0.02% sodium azide in PBS for at least 1 h at room temperature. Retinas were incubated with primary antibodies in PBS containing 3% normal donkey serum, 1% BSA, 0.5% Triton X-100, and 0.02% sodium azide for 5-7 days at room temperature with shaking. Retinas were washed three times in PBS containing 0.5% Triton X-100 (15 min each) before incubation with secondary antibodies in the same diluent overnight at room temperature with shaking. Primary antibodies: rabbit anti-RFP (1:500) and chicken anti-GFP (1:500); secondary antibodies: donkey anti-chicken Alexa Fluor 488 (1:500) and donkey anti-rabbit Cy3 (1:400). Retinas were mounted on glass slides with Dako fluorescent mounting medium.

#### Imaging

Images were acquired on a Nikon-Märzhäuser Slide Express 2 widefield fluorescence microscope, using the following settings: 10x objective (Plan Apo λ, numerical aperture 0.45, optical resolution 0.7-0.8 µm), 16-bit depth, pixel size 0.64 µm. Whole-section and whole-retina images were acquired using multiple tiles with 7% overlap and 4 z-steps (15 μm step size). The Extended Depth of Focus (EDF) module combined the z-stacks into a fully focused composite image.

#### Cell counting and atlas registration

Rabies-labeled input neurons were counted manually in ImageJ^37^ using the Multi-point tool, separately for each channel (GFP and tdTomato) corresponding to inputs to medial and lateral SCs neurons. Sections were registered to the Allen Mouse Brain Atlas using the SHARP-Track toolbox^20^ to obtain the coordinates of each labeled neuron and its assigned brain region.

#### Analysis

Because some animals had sufficient labeling for only one region, upper- and lower-field datasets were analyzed independently. Animals with low whole-brain input counts (<500 cells) or substantial labeling in the intermediate and deep SC layers were excluded from further analysis. For wide-field neurons, 7 mice contributed to upper- and 7 to lower-field SCs analysis (6 mice contributed to both). For narrow-field neurons, 6 mice contributed to upper- and 5 to lower-field analysis (4 mice contributed to both). Gad2-Cre data (n = 4) were obtained from a previous study^31^ and reanalyzed here. To allow for comparison, we included all 4 Gad2 animal data despite 17% of putative starter neurons occurring in the intermediate SC.

The divisions of brain regions and areas were based on the Allen Brain Atlas with modifications described previously39. The whole brain was divided into eight parental regions including CTX (consisting of the isocortex and olfactory areas), cortical subplate (sCTX, consisting of the sCTX and cerebral nuclei), hippocampal formation, TH, HYP, MB, HB, and the 261 subareas thereof. Data S1 shows a full list of all areas. All detected GFP-or tdTomato-expressing cells in SCs were excluded from statistical analysis, to ensure no starter cells were counted as input cells. Additionally, areas with input cells in fewer than half of the datasets for each condition were excluded to ensure consistency of inputs across animals (wide-field: <4 of 7 upper-field, <4 of 7 lower-field; narrow-field: <3 of 6 upper-field, <3 of 5 lower-field). The raw data containing the coordinates of each labeled neuron and its corresponding area are available in Data S2.

To identify robust input regions, we applied two parallel filters based on input strength: areas were required to (1) contribute above a threshold proportion of total inputs, and (2) exceed a threshold relative density (defined as the ratio of an area’s input density, i.e. cell counts/volume of area, to the mean density across the brain, i.e. total cell counts excluding SCs/volume of 261 brain areas excluding SCs).

Threshold values for proportion and density were determined using the Kneedle algorithm^40^ applied to the cumulative distribution of each metric. This algorithm identifies the point of maximum curvature in the cumulative distribution, below which inputs may reflect noise or non-specific labeling. Thresholds were computed separately for each group being compared and the mean threshold was applied to both. For comparisons between cell types (wide-field vs. narrow-field), medial and lateral datasets were pooled within each cell type before threshold computation (proportion ≥ 1%, density ≥ 1.7). For comparisons between upper- and lower-field inputs within a cell type, thresholds were computed separately for each and the mean was applied (wide-field: proportion ≥ 1.1%, density ≥ 1.6; narrow-field: proportion ≥ 1%, density ≥ 1.6).

Effect sizes were quantified as log-odds ratios comparing input proportions between upper and lower visual field domains. Positive values indicate bias toward upper-field representations; negative values indicate bias toward lower-field. Confidence intervals (95%) were estimated using bias-corrected and accelerated (BCa) bootstrap (10,000 iterations), resampling animals with replacement within each group. BCa bootstrap corrects for bias and skewness in the sampling distribution and makes no distributional assumptions. Confidence intervals were adjusted for multiple comparisons across brain areas using false coverage rate (FCR) correction, which controls the expected proportion of confidence intervals that fail to cover their true parameters^41^. Complementary statistical inference was performed using permutation tests (10,000 permutations). For each brain area, a null distribution of log-odds ratios was generated by randomly shuffling group assignments. Two-tailed p-values were calculated as the proportion of permutations with absolute log-odds ratios greater than or equal to the observed value, then adjusted using FDR.

To test whether the upper vs lower visual field bias differed between cell types, we computed the interaction effect as the difference in log-odds ratios between wide- and narrow-field neurons. Positive values indicate stronger upper field bias in wide-field inputs; negative values indicate stronger bias in narrow-field inputs. Bootstrap confidence intervals were estimated by resampling animals with replacement within each of the four group × cell type combinations. For permutation tests, cell type labels were shuffled within each visual field group separately, preserving the main effect of visual field while generating a null distribution for the interaction. Confidence intervals and p-values were adjusted using FCR and FDR, respectively.

Pearson’s correlations were calculated on log-transformed input percentages to reduce the dominance of large projections and linearize the data. To estimate confidence intervals, we used bootstrap resampling (10,000 iterations): for each iteration, animals were resampled with replacement separately within each group, mean log-transformed values were computed across resampled animals, and the correlation was calculated. The 95% confidence interval was defined as the 2.5^th^ and 97.5^th^ percentiles of the bootstrap distribution.

Hierarchical clustering was performed on pairwise Pearson correlations between input areas, computed from input proportions across animals. Clustering used average linkage on correlation distance (1 − r), with cluster boundaries determined at 62% of maximum linkage height to capture visually distinct groups. Similarity between wide-field and narrow-field correlation matrices was assessed using a Mantel test^42^. The observed correlation between matrices was compared to a null distribution generated by permuting area labels (10,000 permutations). The p-value was calculated as the proportion of permuted correlations greater than or equal to the observed value.

To quantify the spatial segregation of neurons projecting to upper versus lower visual field representations within each input region, we identified the axis of maximum separation between the two populations in 3D space. For each animal with dual-color labeling, we projected neuron coordinates of each area onto a set of candidate directions uniformly distributed across a unit sphere. These directions were generated using spherical coordinates, with 36 polar angles (θ) linearly spaced from 0 to π and 72 azimuthal angles (ϕ) linearly spaced from 0 to 2π, yielding 2592 candidate directions with approximately 5° angular spacing. For each candidate direction, we computed kernel density estimates for the upper- and lower-field projecting populations and calculated their overlap. To ensure stable density estimates, only areas with at least 30 neurons per population were included. The direction yielding the minimum overlap was selected as the optimal separation axis. Separation was quantified as 1 minus the overlap integral of the two density estimates, yielding a value ranging from 0 (complete overlap) to 1 (complete segregation). This procedure was performed independently for each animal, and separation values were averaged across animals.

Bar graphs are expressed as median ± median absolute deviation; circles denote individual animals. Density estimates are Gaussian kernel density estimates. 3D renderings of whole-brain input neurons were created using the Python package brainrender^43^. All other analyses were performed using custom MATLAB code.

### MAPseq projection mapping

#### Sindbis virus injections

To trace axonal projections from SCs neurons to downstream targets, we used MAPseq, which labels individual neurons with unique RNA barcodes that are transported to axon terminals and can be detected by sequencing. Mice were injected with a Sindbis viral barcode library (3 × 10^10^ genome copies/ml, diversity of 2 × 10^7^ barcodes; Cold Spring Harbor Laboratories) into the left SCs. Injections were made at 13 sites spanning the medial-lateral and anterior-posterior SCs (75 nl per site; coordinates relative to lambda in mm, AP/ML/DV: +0.2/+0.3/1.30, +0.6/+0.3/1.40, +1.0/+0.3/1.45, +1.3/+0.3/1.45, +0.2/+0.6/1.35, +0.6/+0.6/1.45, +1.0/+0.6/1.50, +1.3/+0.6/1.50, +0.2/+1.0/1.45, +0.6/+1.0/1.60, +1.0/+1.0/1.65, +1.3/+1.0/1.65, −0.1/+0.6/1.20). The titer and volume were similar to previous studies in which single-cell analysis indicated that most neurons express a single barcode, with only a small fraction expressing multiple barcodes^23^. While expression of multiple barcodes within single neurons would lead to overestimation of neuron counts, it does not affect the relative abundance of projection motifs or bulk connection strengths^23,44^.

#### Tissue dissection

After 44–46 hours, mice were transcardially perfused with 4% paraformaldehyde prepared with RNase-free water. Brains were sectioned at 250 μm on a cryostat set to −15°C. All surfaces, including the cryostat, were cleaned with RNase AWAY or 100% ethanol. For each section, the blade was moved to an unused portion and replaced once fully used. Brushes used to collect sections were cleaned with 100% ethanol between sections. Sections were immediately mounted onto slides and placed on dry ice. Target regions were manually dissected on a metal surface surrounded by dry ice to keep sections frozen. Scalpel handles were cleaned with RNase AWAY between sections, and a new blade was used for each target site. For sections containing LP or LG, the optic tract was first removed with a separate blade to minimize contamination from fibers of passage. Source sites collected from the SCs included: SCs-aam, SCs-am, SCs-mm (grouped as medial SCs, corresponding to upper visual field), and SCs-aal, SCs-al, SCs-ml (grouped as lateral SCs, corresponding to lower visual field), as well as SCs-c. Target sites included: contralateral SCs (aam, am, mm, aal, al, ml, c), ipsilateral and contralateral SCd (aam, am, mm, aal, al, ml, c), and ipsilateral and contralateral LP, PBG, LGd, LGv, and MLR. Samples from up to 4 other regions were collected from each brain as negative controls. Dissected tissue was collected on dry ice and stored at −80°C until shipping. Samples were shipped on dry ice to Cold Spring Harbor Laboratories for further processing.

#### Barcode sequencing

Barcode RNA was extracted, reverse transcribed, and amplified using a published protocol^23^. All preprocessing was performed at the MAPseq Core Facility at Cold Spring Harbor Laboratories as described previously (Supplementary Note 4 in ref ^23^), with sequencing performed blinded to sample identity. Briefly, Illumina paired-end reads containing the 30-nucleotide barcode, 2-nucleotide pyrimidine anchor, 12-nucleotide unique molecular identifier (UMI), and 8-nucleotide slice-specific identifier (SSI) were merged into a single file. Reads were demultiplexed by SSI using fastx_barcode_splitter, filtered to remove ambiguous bases, collapsed to unique sequences, and sorted. A minimum read threshold was manually selected to remove low-abundance sequences likely representing PCR or sequencing errors. UMI sequences were removed to convert reads into counts. Spike-in molecules (24-nucleotide barcodes followed by ATCAGTCA) were processed separately. Error correction was performed using bowtie, aligning barcode sequences with up to 3 mismatches. Low-complexity sequences containing stretches of more than six identical nucleotides were removed.

Raw barcode counts were normalized by spike-in RNA abundance and organized into an N × R matrix, where N is the number of detected barcodes (a proxy for neuron count) and R is the number of target regions. Each row represents a single neuron, with values indicating barcode counts (projection strength) in each region. Matrices from all mice were concatenated. Rows containing only zeros were removed to restrict analysis to neurons projecting to at least one target. Data were further filtered to retain only ‘high-confidence’ cell bodies, i.e. barcodes with at least 30 counts and for which the highest barcode count (assigned soma location) was at least tenfold greater than the count in any single target region. Barcodes detected in the negative control region were excluded. A neuron was considered to project to a given target if its barcode count in that region exceeded 2, based on the maximum count observed in negative control samples.

#### Analysis

Because only 3 mice were analyzed, statistical inference was based on neurons pooled across animals unless otherwise stated, allowing valid fixed-effect conclusions on the sampled population of neurons rather than the population of animals^35^. Per-mouse projection strength profiles are shown in Fig. S4a to allow assessment of inter-animal consistency.

To visualize the projection patterns of individual neurons, barcode count matrices were normalized so that each neuron’s counts summed to 1. Neurons were sorted by dominant target, with single-target projections grouped first, followed by multi-target projections sorted by descending projection strength. Sorted matrices were displayed as heatmaps, with white indicating zero projection strength.

Projection probability was defined as the fraction of neurons with non-zero barcode counts in a given target. Differences between upper and lower visual field populations were assessed by bootstrap (10,000 iterations, resampling neurons with replacement within each population). Log_2_ fold change and absolute probability difference were derived from each bootstrap sample. Confidence intervals were corrected for multiple comparisons across targets using FCR adjustment. A difference was considered statistically significant when the corrected confidence interval excluded zero.

Innervation strength was compared between populations among projecting neurons only (those with non-zero barcode counts in a given target). Log_2_ fold changes were estimated by bootstrap (10,000 iterations, resampling with replacement), with confidence intervals corrected using FCR adjustment. Average projection strength per neuron, combining the probability of projecting and the strength of innervation, was computed as mean barcode counts including non-projecting neurons. Log_2_ fold changes were estimated by bootstrap as above.

Normalized projection strength was computed as the fraction of total output directed to each target, i.e. the relative distribution of projections across targets.

To characterize co-projections of SCs output neurons, we computed pairwise conditional projection matrices. For each pair of target areas A and B, we calculated the fraction of upper and lower visual field neurons projecting to area A that also projected to area B.

To identify non-random projection motifs, we compared observed motif frequencies to expected probabilities derived from a binomial model assuming independent projection probabilities. For each ipsilateral and contralateral target, the probability of projection was computed as the fraction of neurons with non-zero barcode counts in that target. Expected motif probabilities were then calculated as the product of probabilities for targets included in the motif and (1 -probability) for targets excluded, assuming independence across targets. All possible motifs involving one to six targets were enumerated, and expected counts were computed by multiplying expected probabilities by the total number of neurons. For each motif, we tested whether observed counts deviated from expected using a two-tailed binomial test, with p-values corrected for multiple comparisons using FDR. To compare motif distributions between upper and lower visual field populations, we computed the difference between observed and expected motif probabilities for each population. Cumulative distributions of these differences were compared using a two-sample Kolmogorov-Smirnov test. For individual motifs, differences in observed probabilities between populations were assessed using Fisher’s exact test with FDR correction.

### Visual responses of wide-field neurons

#### Labeling of wide-field neurons and cranial window implantation

Wide-field neurons were labeled using adeno-associated viral (AAV) injections in Ntsr1-GN209-Cre mice (8– 14 weeks old) combined with cranial window implantation over the superior colliculus.

Mice were anesthetized with ketamine (75 mg/kg) and medetomidine (1.0 mg/kg, intramuscular) and placed on a heating pad maintained at 37°C. The head was stabilized in a stereotaxic frame using ear bars. After scalp removal, a titanium headplate with an integrated ring was fixed to the skull (centered 2 mm lateral of midline and 2 mm anterior of lambda) using super glue and dental cement. Animals were subsequently head-fixed via the implanted headplate.

A 4 mm craniotomy was performed over the superior colliculus. The dura was removed, and overlying cortex was aspirated under continuous application of artificial cerebrospinal fluid (ACSF) to prevent desiccation and control bleeding. Viral injections (AAV1.Syn.Flex.GCaMP6s.WPRE.SV40) were performed using glass micropipettes (10 μm inner diameter). Up to six sites were targeted based on the retinotopic organization of the superior colliculus. At each site, 200 nL of virus was injected at depths of 100, 200, and 400 μm, with 3-minute intervals between injections.

Following injections, a cylindrical glass window was placed over the exposed superior colliculus and secured with tissue adhesive. A custom rubber ring was mounted on the headplate to form a reservoir for two-photon imaging and sealed with adhesive. The implant was reinforced with dental cement.

Postoperatively, mice received buprenorphine (0.2 mg/kg, i.p.) and cefazolin (15 mg/kg, i.p.) every 12 h for 72 h, and body weight was monitored during recovery.

#### In vivo two-photon calcium imaging

Calcium activity of wide-field neurons in the superficial superior colliculus was recorded using a two-photon microscope (Thorlabs B-Scope) with a Ti:Sapphire laser (920 nm) to excite GCaMP6s. Imaging was performed at 30 Hz using Scan-Image 4.2. During recordings, mice were awake and head-fixed on an air-supported spherical treadmill.

Visual stimuli were presented in the upper (elevation > 0°, n=4) or lower visual field (elevation ≤ 0°, n=6) and were warped to correct for projection onto a curved dome surface. Stimuli consisted of either a 2° black or white disc linearly expanding to 45° within 1s, a 45° black disc linearly shrinking to 2° or a 45° grey disc linearly dimming to black within 1s, each repeated 10 times. Sweeping black or white discs of 2°, 8°, or 32° traversed 240° of visual space at constant elevation at either 30°/s or 160°/s and in two directions, with each condition repeated three times. Optimal stimulus position was determined by presenting a 10° flashed black disc at 25 locations (combinations of −20°, 0°, 20°, 45°, and 90° azimuth with −20°, 0°, 20°, 40°, and 60° elevation). The location eliciting the strongest response in the recording area was selected.

#### Analysis

Two-photon imaging data was preprocessed using the CaImAn pipeline^45^ for motion correction and region-of-interest (ROI) detection. Neurons were manually identified from the detected ROIs and calcium traces extracted as ΔF/F using custom scripts in Python 3.7.

A neuron’s responsiveness to a particular stimulus was assessed by calculating the quality index (QI)^46^, which measures trial-to-trial response consistency. Based on the bimodal distribution of QI values, neurons with QI > 0.2 were classified as responsive to a stimulus.

Statistical analysis was performed using bootstrap resampling (10,000 iterations) on neurons pooled across animals. Differences between means correspond to a permutation test (non-parametric two-sample test), without assuming normality. Statistical significance was defined as p < 0.05.

### Disynaptic output tracing of wide-field neurons

#### Transsynaptic tracing injections

For transsynaptic output tracing, two adeno-associated viral (AAV) constructs were used. First, 50 nl of AAV1-hSyn-Cre-WPRE-hGH (Addgene #105553) was injected to drive expression of Cre recombinase under the human synapsin promoter. Second, 50 nl of AAV1-phSyn1(S)-flex-tdTomato-T2A-SypEGFP-WPRE (Addgene #51509) was used as a Cre-dependent reporter to drive expression of cytoplasmic tdTomato and synaptophysin-tagged GFP under the synapsin promoter.

The first injection was targeted to either the medial (n=4) or lateral (n=1) SCs of C57BL/6J wild-type mice, resulting in Cre expression in collicular neurons. Viral particles were transported anterogradely and spread transsynaptically to neurons in the lateral posterior nucleus (LP), as previously described^47^. One week later, the second AAV was injected into the LP. Neurons transduced by both constructs expressed tdTomato throughout the cytoplasm and EGFP at presynaptic terminals, enabling visualization of cell bodies and axonal projections.

Stereotaxic coordinates were determined using the Allen Mouse Brain Atlas for anterior-posterior (AP) and medio–lateral (ML) positioning, and the Paxinos and Franklin Mouse Brain Atlas for dorsal–ventral (DV) coordinates. Injection coordinates were as follows:

Medial SCs: AP −3.7 mm; ML 0.3−0.4 mm; DV −1.20 mm

Lateral SCs: AP −4.0 mm; ML 1.1−1.2 mm; DV −1.25 mm

For LP injections, coordinates depended on the site of the first SCs injection:

Following medial SCs injection: AP −2.1 mm; ML 1.40 mm; DV −2.7 mm

Following lateral SCs injection: AP −2.3 mm; ML 1.40 mm; DV −2.7 mm

#### Brain Immunohistochemistry

Immunofluorescence staining was performed on selected brain sections (n = 6) to enhance fluorescent protein signals and label cell nuclei with DAPI (1:1000). Sections were incubated in blocking buffer (PBS, Triton X-100, 10% donkey serum) for 1 h at room temperature, followed by three washes in PBT. Slices were then incubated overnight at 4°C with primary antibodies diluted in blocking buffer. After washing, sections were incubated with secondary antibodies and DAPI for 4 h at room temperature in the dark. Finally, sections were washed, mounted on glass slides, and coverslipped with mounting medium.

Primary antibodies included goat anti-RFP and rabbit anti-GFP (both 1:1000). Corresponding secondary antibodies were donkey anti-goat Alexa 555 and donkey anti-rabbit Alexa 488 (both 1:1000). These were used to amplify cytoplasmic tdTomato and synaptic GFP signals, respectively.

#### Imaging

Brain sections were initially screened using a wide-field fluorescence microscope (Olympus MVX10) to verify labeling and identify regions of interest. High-throughput imaging of selected slides was performed using a Nikon-Märzhäuser SlideExpress 2 slide scanner with a 20× objective (0.64 μm/pixel). Images were processed in NIS Elements with extended depth-of-focus and downscaled for analysis. Fluorescent cell bodies were manually counted in ImageJ, and regions of interest were saved for subsequent registration.

For higher-resolution analysis of axon terminals, selected regions were imaged using a confocal microscope (Zeiss LSM 710, 20× Epiplan-Apochromat objective, 0.7 NA) at 0.5 μm/pixel lateral resolution and 0.5 μm in the z-axis. Z-stacks were acquired and processed in Fiji, including channel multiplication to enhance co-localized signals.

Histological images were registered to the Allen Mouse Brain Atlas using the SHARP-Track MATLAB interface^20^.

