## Supplemental Materials for "Visual field position shapes input sampling and output routing in the superior colliculus"

Alex Calzoni *et al*.

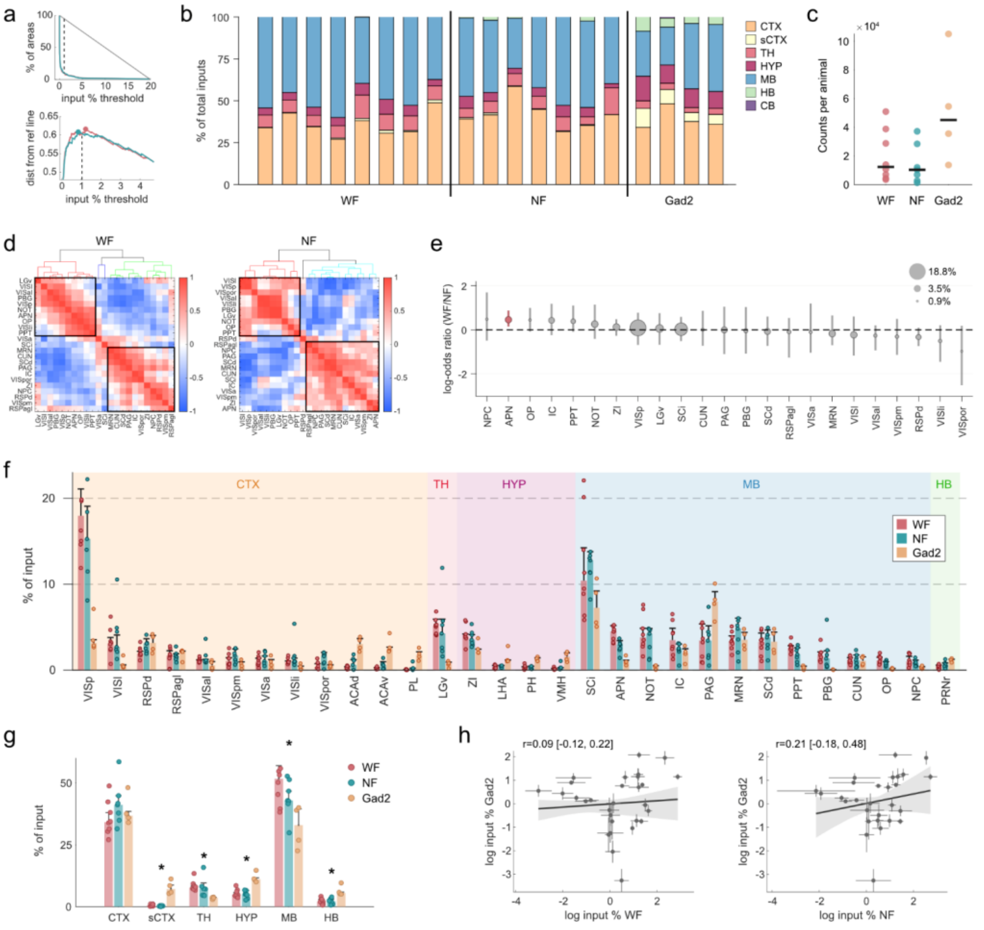

**Fig S1, related to Fig. 1**

**Fig. S1 | Brain-wide inputs to wide-field, narrow-field and Gad2 neurons. a** Kneedle algorithm for identifying input threshold. Top: cumulative distribution of input proportions to wide-field (red) and narrow-field (teal) neurons across brain areas. Bottom: distance from reference line; dashed line indicates average point of maximum curvature (1% threshold). **b** Proportion of inputs by major brain division for individual animals. Each column represents one animal. **c** Total input counts per animal. Black lines denote median. **d** Hierarchical clustering of input areas based on pairwise Pearson correlations of input proportions across animals. Correlation between matrices was assessed by Mantel test ($r = 0.450$, $p = 0.0001$). **e** Log-odds ratios comparing input proportions between wide-field and narrow-field neurons, sorted by effect size. Positive values indicate wide-field bias; negative values indicate narrow-field bias. Error bars show 95% bootstrap confidence intervals with false coverage rate correction. Circle size reflects mean input percentage. **f** Input strength for each brain area as percentage of total inputs. Areas grouped by cortex (CTX), thalamus (TH), hypothalamus (HYP), midbrain (MB), and hindbrain (HB). Bars show median ± MAD; dots represent individual animals. **g** Proportion of inputs by major brain division. Bars show median ± MAD; dots represent individual animals. Asterisks indicate regions where Gad2 inputs differed from both wide- and narrow-field neurons ($p< 0.05$ after FDR correction, Kruskal-Wallis test). **h** Correlation of log-transformed input percentages between wide-field and Gad2 neurons (left) and narrow-field and Gad2 neurons (right). Each point represents one brain area; error bars show ± MAD. Shading indicates 95% bootstrap confidence interval.

**Fig. S2, related to Fig. 2**

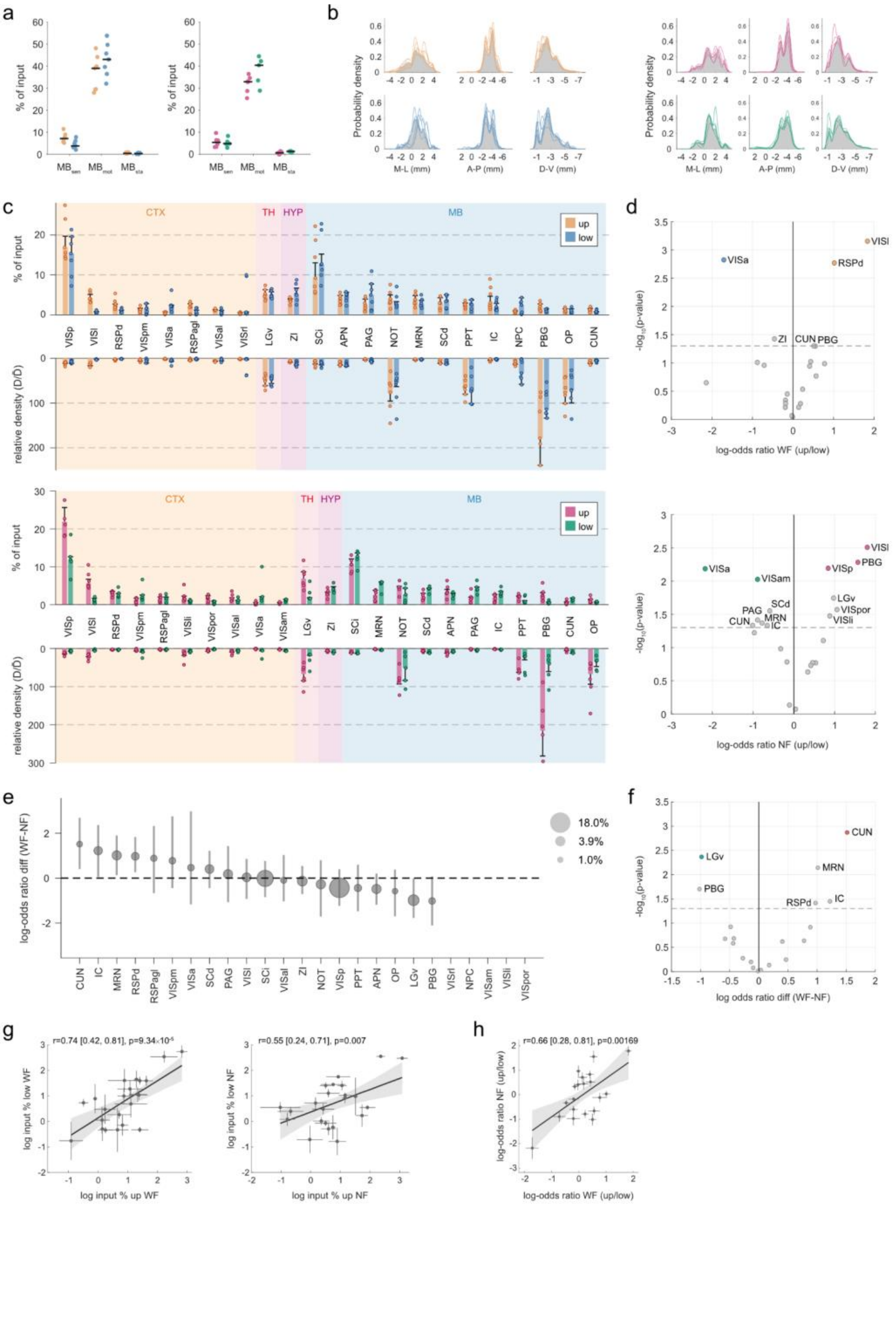

**Fig. S2 | Domain-specific input biases for wide-field and narrow-field neurons.** **a** Input proportions from midbrain subdivisions (MBsen, sensory; MBmot, motor; MBsta, state) for wide-field (left) and narrow-field (right) neurons. Orange/blue denote upper/lower visual field for wide-field neurons; magenta/green denote upper/lower visual field for narrow-field neurons. **b** Probability density distributions of input neuron positions along medial-lateral (M-L), anterior-posterior (A-P), and dorsal-ventral (D-V) axes for wide-field (left) and narrow-field (right) neurons. Gray shading shows the mean density distribution; colored lines represent individual animals. **c** Input strength for each brain area as percentage of total inputs and relative density for wide-field (top) and narrow-field (bottom) neurons. Areas grouped by cortex (CTX), thalamus (TH), hypothalamus (HYP), and midbrain (MB). Bars show median ± median absolute deviation; dots represent individual animals. **d** Volcano plots showing permutation test results for domain bias in wide-field (top) and narrow-field (bottom) neurons. Labeled areas have $p < 0.05$; colored points indicate $p < 0.05$ after FDR correction. **e** Difference in log-odds ratios between wide-field and narrow-field neurons. Error bars show 95% bootstrap confidence intervals with false coverage rate correction. Circle size reflects mean input percentage. **f** Volcano plot showing permutation test results for the difference in log-odds ratios between cell types. Labeled areas have $p < 0.05$; colored points indicate $p < 0.05$ after FDR correction. **g** Correlation of log-transformed input percentages between upper and lower visual field for wide-field (left) and narrow-field (right) neurons. Each point represents one brain area; error bars show ± MAD. Shading indicates 95% bootstrap confidence interval. **h** Correlation of log-odds ratios between wide-field and narrow-field neurons. Each point represents one brain area; error bars show ± MAD. Shading indicates 95% bootstrap confidence interval. Throughout, orange/blue denote upper/lower visual field for wide-field neurons; magenta/green denote upper/lower visual field for narrow-field neurons.

### Fig. S3, related to Fig. 3

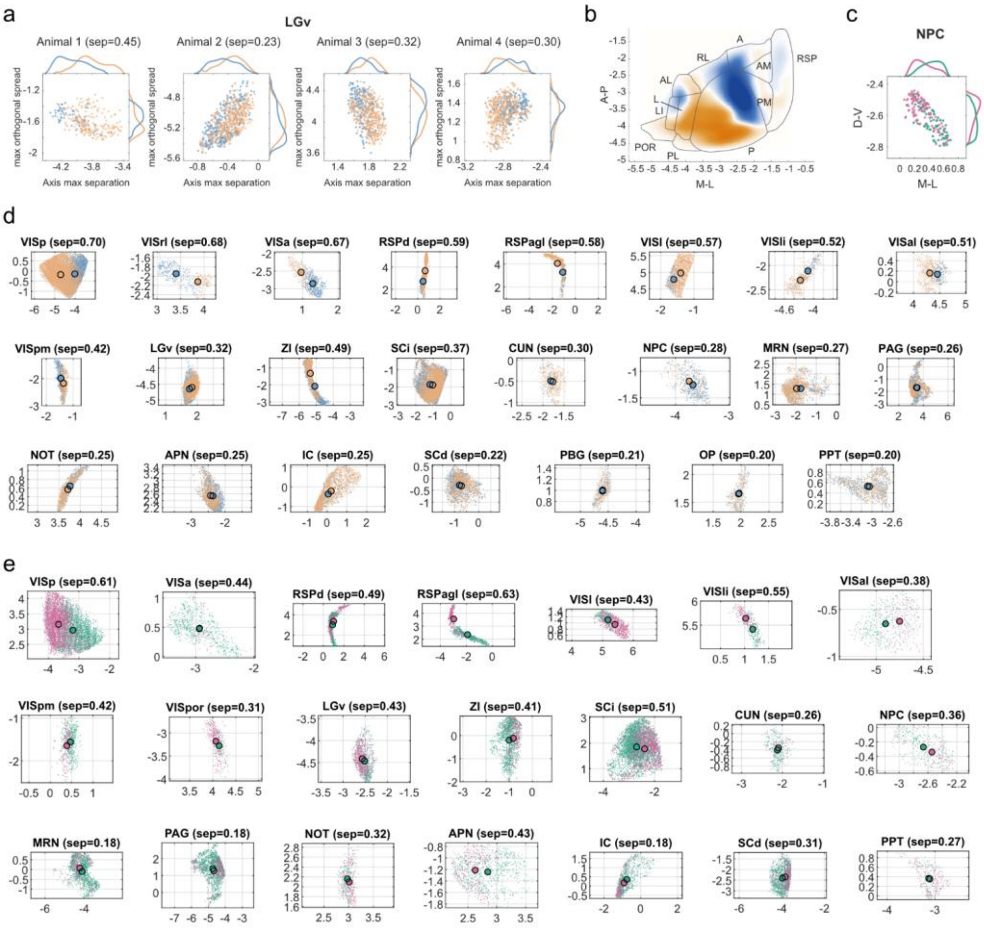

**Fig. S3 | Topographic segregation analysis for individual input regions. a** Example separation analysis for LGv across individual animals. Neurons are projected onto the axis of maximum separation (x-axis) and the axis of maximum orthogonal spread (y-axis, computed via PCA on residuals). Distributions show kernel density estimates along each axis. Separation values indicated per animal; the average across animals is used for the heatmap in Fig. 4a. **b** Kernel density estimates of inputs to wide-field neurons projected onto visual and retrosplenial cortical areas, extending the retinotopic organization shown in Fig. 4c. **c** Nucleus of the posterior commissure showing topographic segregation along the dorsal-ventral axis. **d** Spatial segregation for all input regions to wide-field neurons, sorted by separation value. Each panel shows neurons pooled across animals, projected onto the axis of maximum separation (x-axis) and orthogonal spread (y-axis). Large dots indicate population centroids. Separation values are averaged across individual animals. **e** Same as d for narrow-field neurons. Throughout, orange/blue denote upper/lower visual field for wide-field neurons; magenta/green denote upper/lower visual field for narrow-field neurons.

**Fig. S4, related to Fig. 4**

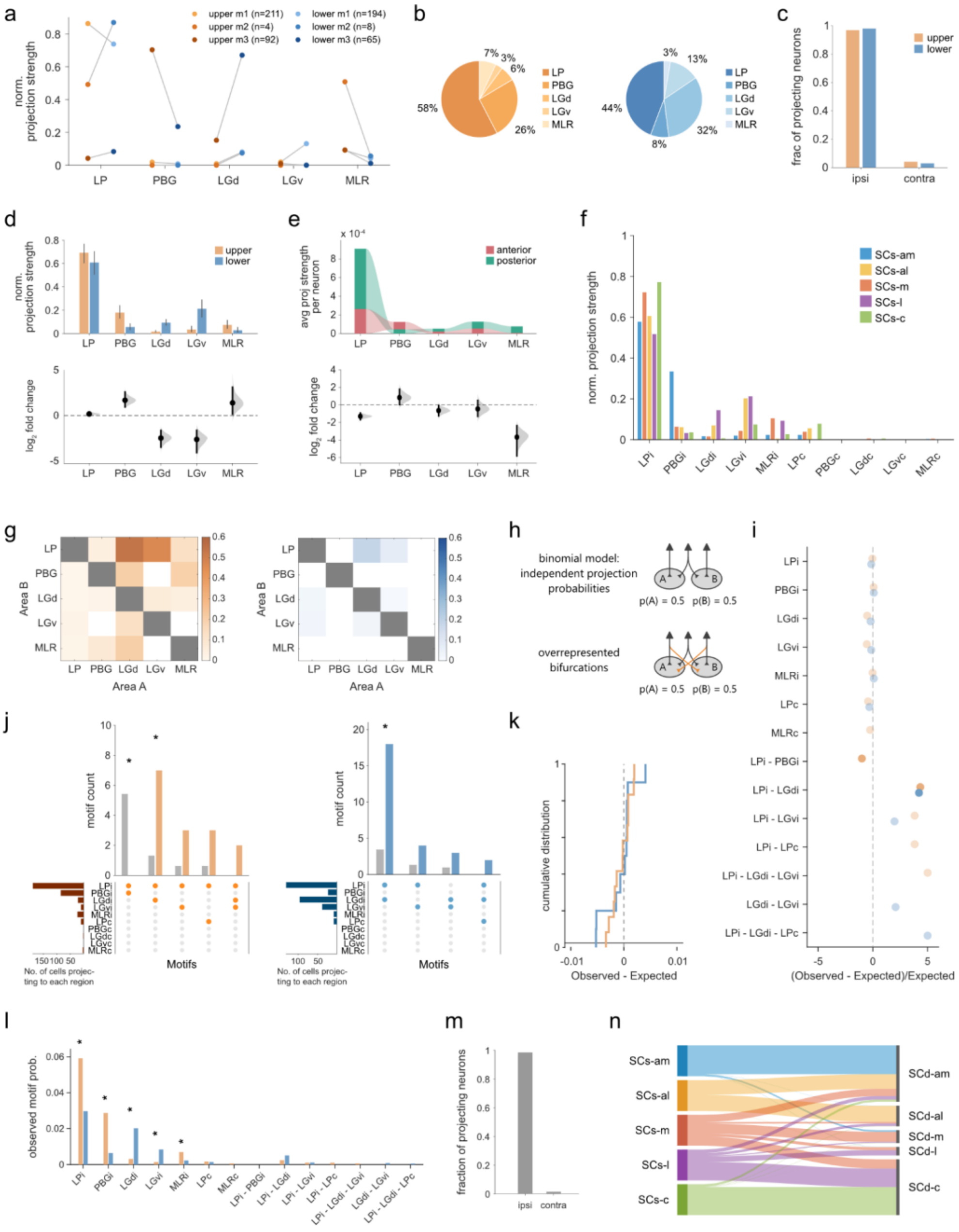

**Fig. S4 | MAPseq projection analysis of collicular neurons. a** Normalized projection strength for individual mice. Individual mice showed consistent direction with the pooled effect for PBG, LGd, and MLR, but not for LGv. LP showed no significant effect in the pooled analysis (see panel d). **b** Proportion of neurons projecting to each target for upper (left) and lower (right) SCs. **c** Fraction of neurons with ipsilateral versus contralateral projections to targets outside the SC. **d** Normalized projection strength (top) and $\log_2$ fold-change (bottom) between upper and lower SCs neurons. **e** Average projection strength per neuron (top) and $\log_2$ fold-change (bottom) between anterior (red) and posterior (green) SCs neurons. Error bars show 95% bootstrap confidence intervals corrected for multiple comparisons (false coverage rate). **f** Normalized projection strength for each SCs source region before grouping into upper and lower populations (am, anteromedial; m, medial; al, anterolateral; l, lateral; c, caudal). For target areas, i denotes ipsilateral, c contralateral. **g** Conditional probability matrices showing the probability of projecting to area B given projection to area A for upper (left) and lower (right) SCs neurons. **h** Binomial model assuming independent projection probabilities (top) and an overrepresented bifurcation motif deviating from the null model (bottom). **i** Normalized deviation from expected motif counts for each projection motif. Dashed line indicates expected value under the binomial model. Faded dots indicate motifs not significantly different from expected. **j** Projection motif counts for upper (left) and lower (right) SCs neurons. Colored bars represent observed counts, gray bars show expected counts under the binomial model. Dots indicate target combinations. Horizontal bars show number of neurons projecting to each region. Asterisks denote motifs significantly over- or underrepresented (binomial test, FDR-corrected). **k** Cumulative distribution of differences between observed and expected motif probabilities. **l** Observed motif probabilities for single-target and multi-target projections. Asterisks denote motifs with significantly different probabilities between upper and lower SCs (Fisher's exact test, FDR-corrected). **m** Fraction of neurons with ipsilateral versus contralateral projections to the SCd. **n** Normalized projection strength from superficial to deeper SC layers (am, anteromedial; m, medial; al, anterolateral; l, lateral; c, caudal). Throughout, orange denotes upper SCs and blue denotes lower SCs.

### Extended results S1, related to Fig. S4

*Projection motifs largely follow independent targeting probabilities*

Conditional probability analysis showed that LGd and LGv innervation was frequently paired with LP projections, especially for upper SCs neurons (Fig. S4g). We therefore investigated whether SCs neurons exhibit non-random projection motifs. We compared observed projection motifs to expected probabilities from a binomial model assuming independent projection probabilities (Fig. S4h).

We identified one underrepresented and two overrepresented projection motifs for upper SCs neurons, and one overrepresented motif for lower SCs neurons (Fig. S4j). The only underrepresented motif was the bifurcation to ipsilateral LP and PBG, which was not observed among upper SCs neurons despite an expected count of 5.5 ($p= 0.002$). This reflects PBG receiving the highest proportion of dedicated projections (Fig. 5d). Among overrepresented motifs, the LP-LGd bifurcation was shared between medial and lateral neurons, whereas the bifurcation to ipsi- and contralateral LP occurred only in medial neurons.

Although multi-target motifs frequently involved LP, LGd, and LGv, overall counts were low because most neurons projected to single targets. In fact, most motifs could be explained by the binomial model (Fig. S4i). In support of this, the cumulative distributions of differences between observed and expected motif probabilities did not differ between upper and lower visual field (two-sample Kolmogorov-Smirnov test, $p=0.51$; Fig. S4k). Deviations from independence were small for both populations, indicating that individual neurons select their projection targets largely independently of one another regardless of retinotopic position. This apparent contradiction, high single-target specificity despite assumed independence, is explained by the intrinsically low probability of projecting to any individual target (Fig. S4l).

*Superficial SCs subdivisions preferentially target their corresponding deeper layers*

We analyzed local connectivity from superficial to deeper SC layers. Because ipsilateral projections account for 98.5% of SC output (Fig. S4m), we restricted analysis to the ipsilateral hemisphere to simplify interpretation. Each SCs subdivision (SCs-am, SCs-al, SCs-m, SCs-l, SCs-c) predominantly targeted its corresponding deeper region (n=5,158, Fig. S4n), consistent with established topographic organization where superficial layers relay visual input to functionally aligned deeper layers involved in multisensory integration.

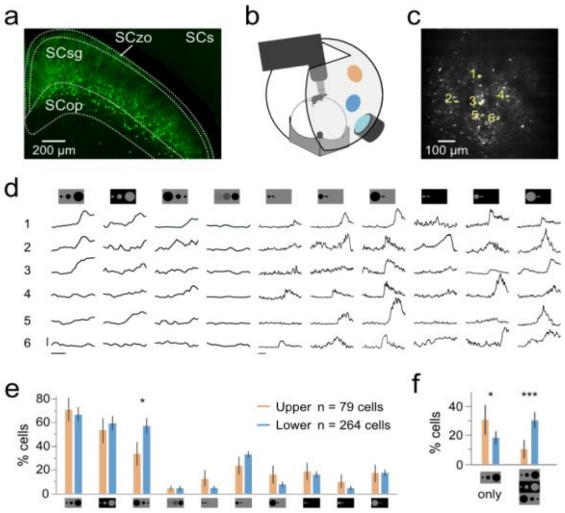


**Fig. S5 | Wide-field neurons show visual field-dependent differences in stimulus selectivity. a** Virally labeled wide-field neurons in SCs. **b** Setup for headfixed two-photon calcium imaging with presentation of upper- and lower-field stimuli in a dome. **c** Imaging area with highlighted example neurons. **d** Example responses of upper- and lower-field wide-field neurons to black and white looming, black shrinking, dimming and black and white sweeping discs. **e** Percent responding wide-field neurons to stimuli in d. Upper 79 cells, n = 4, lower 264 cells, n = 6. *p = 0.017 (two-sample permutation test). **f** Percent responding wide-field neurons to exclusively black looming vs black and white looming and shrinking discs. Upper 61 cells, n = 4, lower 226 cells, n = 6. *p = 0.013, ***p < 0.001 (two-sample permutation test). Throughout, orange denotes upper and blue lower field. Bars indicate mean and error bars standard deviation from bootstrapping.

### Extended results S2, related to Fig. S5

*Upper- and lower-field wide-field neurons have distinct visual feature* selectivity

Differences in output routing raised the possibility that upper- and lower-field modules also differ in the visual signals they carry. To test this in a defined cell type, we recorded calcium responses from virally labeled wide-field neurons in head-fixed mice running on an air-cushioned ball while visual stimuli were presented on a spherical screen (Fig. S5a,b). Responses to black and white looming discs, black and white sweeping discs, shrinking black discs, and dimming black discs were measured at upper- and lower-field locations (Fig. S5c). Because responses were pooled across animals, the resulting comparisons apply to the sampled neuron populations. Upper- and lower-field wide-field neurons differed in their response profiles within this stimulus set. Lower-field neurons showed a higher response fraction to shrinking discs (Fig. S5d). Upper-field neurons, by contrast, were more likely to respond selectively to black looming discs, whereas lower-field neurons more often responded to multiple related stimulus classes, including black and white looming and shrinking discs (Fig. S5e). These data indicate that wide-field neurons in upper- and lower-field SC domains differ in stimulus selectivity within the tested stimulus space.

**Commented [KF1]:** I'd make this a smaller square with traces as D set below a,b,c then d and e as final row.

**Commented [AC2R1]:** check to see if it's what you envisioned

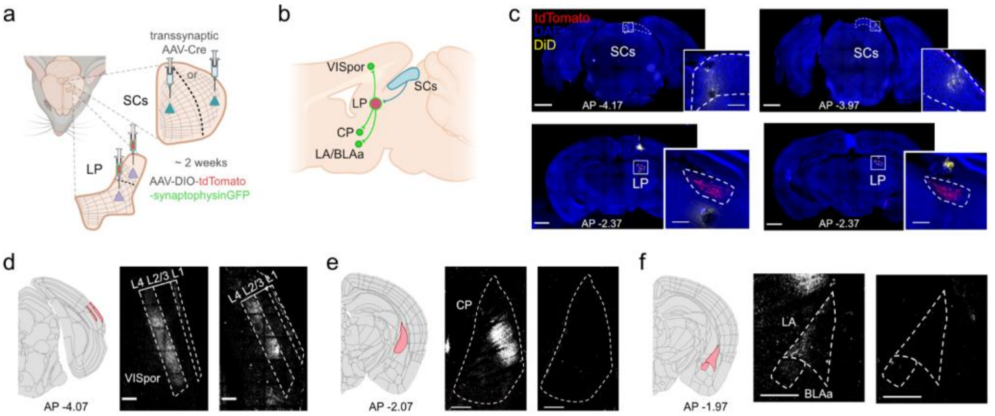


**Fig. S6 | Disynaptic tracing through LP suggests partially distinct downstream routing. a** Transsynaptic tracing strategy of disynaptic projection targets of upper (left) and lower (right) wide-field neurons. **b** Primary (red) and disynaptic target regions (green). **c** Coronal sections of upper (left) and lower (right) injections in SCs (top) and LP (bottom). Scale bars 1000 μm, 200 μm in insets. **d-f** Coronal sections of disynaptic projections in VISpor (d, layers 1, 2/3 and 4 indicated), CP (e) and LA and BLAa (f). Area and AP location (left), GFP/tdTomato colabeled axonal boutons of disynaptic projections of upper (middle) and lower (right) wide-field neurons indicated in white. Scale bar 500 μm.

### Extended results S3, related to Fig. S6

*Disynaptic tracing through LP suggests partially distinct downstream routing*

Differences in wide-field stimulus selectivity raised the possibility that upper- and lower-field circuits also differ in the downstream pathways they access through LP. To test this, we used transsynaptic AAV1-Cre injected into medial or lateral SCs together with a Cre-dependent reporter in LP to label secondary projection targets of wide-field neurons (Fig. S6a-c). Secondary targets were identified in coronal sections by co-labeling of cytosolic and axonal markers. Both upper- and lower-field wide-field circuits reached postrhinal cortex (VISpor; Fig. S6d). In the present dataset, labeling in caudoputamen and amygdala was observed only from the upper-field condition (Fig. S6e,f). Because the lower-field condition was represented by a single animal, we interpret this difference cautiously.

**Commented [KF3]:** I'd make this two rows. As compact as possible.

**Data S1, related to Fig. 2-4.**

Sheet "Brain areas combined list" indicates details of the 261 subareas used in the analysis, abbreviations ("Acronym"), full name ("Full Name"), major regions belonging to ("Parental Area"), and the combined Allen mouse brain atlas indexes of the brain areas ("Combined Area Idx").

Sheet "Allen mouse brain areas" contains all the areas from the Allen mouse brain atlas (a reference to the "Combined Area Idx").

**Data S2, related to Fig. 2-4.**

Brain area ("name" and "acronym") and coordinates (from bregma in mm: anterior-posterior: "AP_location", dorso-ventral: "DV_location" and medio-lateral: "ML_location") of all counted cells. Column "avIndex" indicates the area identifier used in the Allen mouse bran atlas. Each sheet contains cells of individual mice.
